# Aerial High-Throughput Phenotyping Enabling Indirect Selection for Grain Yield at the Early-generation Seed-limited Stages in Breeding Programs

**DOI:** 10.1101/2020.04.21.054163

**Authors:** Margaret R. Krause, Suchismita Mondal, José Crossa, Ravi P. Singh, Francisco Pinto, Atena Haghighattalab, Sandesh Shrestha, Jessica Rutkoski, Michael A. Gore, Mark E. Sorrells, Jesse Poland

## Abstract

Breeding programs for wheat and many other crops require one or more generations of seed increase before replicated yield trials can be sown. Extensive phenotyping at this stage of the breeding cycle is challenging due to the small plot size and large number of lines under evaluation. Therefore, breeders typically rely on visual selection of small, unreplicated seed increase plots for the promotion of breeding lines to replicated yield trials. With the development of aerial high-throughput phenotyping technologies, breeders now have the ability to rapidly phenotype thousands of breeding lines for traits that may be useful for indirect selection of grain yield. We evaluated early generation material in the irrigated bread wheat (*Triticum aestivum* L.) breeding program at the International Maize and Wheat Improvement Center to determine if aerial measurements of vegetation indices assessed on small, unreplicated plots were predictive of grain yield. To test this approach, two sets of 1,008 breeding lines were sown both as replicated yield trials and as small, unreplicated plots during two breeding cycles. Vegetation indices collected with an unmanned aerial vehicle in the small plots were observed to be heritable and moderately correlated with grain yield assessed in replicated yield trials. Furthermore, vegetation indices were more predictive of grain yield than univariate genomic selection, while multi-trait genomic selection approaches that combined genomic information with the aerial phenotypes were found to have the highest predictive abilities overall. A related experiment showed that selection approaches for grain yield based on vegetation indices could be more effective than visual selection; however, selection on the vegetation indices alone would have also driven a directional response in phenology due to confounding between those traits. A restricted selection index was proposed for improving grain yield without affecting the distribution of phenology in the breeding population. The results of these experiments provide a promising outlook for the use of aerial high-throughput phenotyping traits to improve selection at the early-generation seed-limited stage of wheat breeding programs.

## INTRODUCTION

For small grains as well as many other crops such as soybean and rice, limited seed yield per plant after deriving fixed lines can delay replicated yield testing until sufficient quantities of seed are accumulated through one or more generations of seed increase. As a result, breeding lines are sown in rows or small plots in the field for the purpose of seed increase prior to yield testing. The generation at which this occurs depends on the crop and breeding scheme followed, though in theory this may begin as early as the F_3_ with yield testing at the F_4_. Breeding lines at this stage are also typically subject to visual selection, largely for phenology and disease resistance, as more extensive phenotyping is limited by time and costs. The ability to accurately cull breeding lines with low potential in the early generation seed-limited stage is highly desirable because it reduces subsequent expenditure on costly yield testing (Brennan, 1988).

Selection at the early generation seed-limited stage of a breeding program relies on the assumption that the phenotype of a breeding line when sown in a small, unreplicated plot is predictive of its eventual performance in a larger replicated yield trial. Selection on small plots of unreplicated entries can be applied most effectively for qualitative traits controlled by a limited number of genetic loci with major effects, high heritability, and limited genotype-by-environment interaction (Snape & Simpson, 1984). Selection on quantitative traits, however, becomes more challenging due to lower heritability and difficulty in obtaining effective quantitative measurements (Bernardo, 2003).

Numerous studies have shown that plants exhibit plastic phenotypic responses to the competitive stress imposed by their neighbors (White & Harper, 1970; Edmeades & Daynard, 1979; Dornbusch et al., 2011). The level of competitiveness occurring in small plots does not completely mimic that which occurs in yield trials or farmers’ fields, though it will be more representative than single space-planted plants. In addition, the small plot size increases error in grain yield measurements. The yields of single wheat plants or small plots have been observed to exhibit low heritability (Fonseca & Patterson, 1968; Mitchell et al., 1982) and weak correlations with the yields of replicated yield trials (Syme, 1972; Knott & Kumar, 1975). Non-genetic factors can also drive variation in trait values when breeding material is evaluated in small plots (reviewed by Rebetzke et al., 2014). For example, variation in canopy height and architecture can lead to light competitiveness among different breeding lines sown in adjacent plots (Clarke et al., 1998). Finally, the large number of breeding lines at this stage of the program is effectively prohibitive for these desired measurements of grain yield. Therefore, it can be difficult to directly and accurately select for grain yield at the early generation seed-limited stages of breeding programs.

A potential strategy to improve selection accuracy at the early generation seed-limited stages of wheat breeding programs would be to perform indirect selection on a highly heritable secondary trait, namely traits that exhibit strong correlations with grain yield when measured in replicated yield trials (reviewed by Fischer & Rebetzke, 2018). Several measurable physiology traits in wheat have been shown to fit these criteria. For example, Rebetzke et al. (2002) demonstrated that selection for reduced carbon isotope discrimination at the early generation stage increased grain yield and aboveground biomass in wheat when grown under drought stress. Quail et al. (1989) found significant correlations between wheat grain yield in the F_7_/F_8_ generation with fruiting efficiency at maturity when measured in the F_3_ generation. In Condon et al., (2008) stomatal aperture-related traits measured on 1.6 m^2^ plots in the F_4_-F_6_ generations were as effective as visual selection in identifying wheat breeding lines with high yield potential.

While these and other studies demonstrate the potential for improving grain yield through indirect selection on secondary traits, the feasibility of applying these approaches within breeding programs remains limited due to the high cost and time required to evaluate these traits in the field on large numbers of breeding lines. For indirect selection at the early generation seed-limited stage to be feasible, the phenotyping approach must be amendable to high-throughput data collection to cover the large number of breeding lines sown, and the cost must not exceed the amount saved on subsequent yield trials by culling low potential lines based on secondary traits.

Recent applications of advances in the remote sensing industry to plant breeding have aimed to enable rapid, field-based phenotyping of physiological and other traits on large numbers of breeding lines while reducing labor, time, and cost with respect to traditional approaches (Cabrera-Bosquet et al., 2012; Araus & Cairns 2014; Pauli et al., 2016). In recent years, a range of ground- and aerial-based high-throughput phenotyping (HTP) systems utilizing various cameras and sensors have been developed and tested to expand the efficiency of phenotypic data collection (Andrade-Sanchez et al., 2014; Crain et al., 2016; Haghighattalab et al., 2016; Watanabe et al., 2017). These proof-of-concept approaches have now reached a validation stage to be applied in breeding programs for improved selections.

Vegetation indices (VIs), which provide an integrated measurement of canopy structure and photosynthetic activity based on the amount of light reflected off of the crop canopy (Huete et al., 2000; Reynolds & Langridge, 2016), are highly amendable to HTP. A number of studies have demonstrated strong correlations between VIs and grain yield in wheat (Babar et al., 2006; Prasad et al., 2007; Prasanna et al., 2013). In addition, spectral reflectance traits have been shown to improve the accuracy of grain yield prediction in wheat when integrated with genetic information in genomic selection (GS) (Rutkoski et al., 2016; Sun et al., 2017; Crain et al., 2018; Krause et al., 2019). These previous efforts have leveraged the replicated yield trial stage of breeding programs to evaluate the ability to predict grain yield using traits collected with aerial HTP systems. While these approaches may serve to provide breeders with yield predictions earlier in the season, thereby facilitating greater resource-use efficiency at harvest, the potential cost savings may not justify the expense of collecting the HTP traits. Furthermore, once a field experiment with full-size, replicated plots has been established, it is in the best interest to complete harvesting for a direct measure of grain yield. In the pipeline of a breeding program, a greater benefit could come from applying HTP at the early stages of the breeding cycle when limited seed availability prevents the assessment of grain yield in replicated full-sized plots. If HTP traits can facilitate greater selection accuracy at early stages, programs can select at a higher intensity, thereby promoting fewer and better lines to more expensive replicated yield trials.

Plant breeding strategies in small grains – including pedigree selection, bulk selection, single seed descent, and doubled haploids, among others – require one or more generations of spaced plantings for seed increase prior to replicated yield testing, opening up the possibility for indirect selection on secondary traits collected with HTP. The bread wheat breeding program at the International Maize and Wheat Improvement Center (CIMMYT) utilizes a “selected bulk” scheme from the F_2_ through F_5_ generations in which selected spikes from a cross are harvested and bulked to form the next generation (Singh et al., 1998). At the F_5_ or F_6_ generation, individual plants or spikes are selected and promoted to the next stage as independent derived lines, which are sown in beds as double-row 0.7m × 0.8m unreplicated small plots (SP) at the Campo Experimental Norman E. Borlaug (CENEB) in Ciudad Obregón, Sonora, México. This method enables tens-of-thousands of breeding lines to be planted within a small area for seed increase and visual selection prior to the first year of replicated yield testing. In the field, lines are selected and advanced based on uniformity, disease resistance, and other agronomic characteristics. A second round of visual selection is then performed on the seed following harvest to retain breeding lines with desirable grain characteristics for promotion to the first year of replicated yield trials (YT) (van Ginkel et al., 2002).

Detailed phenotyping of the SP stage of CIMMYT’s irrigated bread wheat breeding program is challenging due to the large number of lines sown each breeding cycle. As an example, over 40,000 lines were sown as SP at CENEB during the 2016-17 cycle. Aerial-based HTP platforms, however, enable screening of thousands to tens-of-thousands of plots within a short time span and with less labor, opening up the possibility of selection on secondary HTP traits at the SP stage of the CIMMYT wheat breeding program.

A potential challenge that may limit the utility of aerial HTP for use in indirect selection for grain yield is the strong associations between VIs and phenology that are often observed (Magney et al., 2016; Duan et al., 2017). When this confounding of traits occurs, selection schemes based on the relationship between VIs and grain yield would indirectly affect the distribution of phenology, disproportionally favoring later maturing entries. Although late maturity often confers superior yields, it is not always favorable as it may expose the crop to terminal and continual high temperature stress and places constraints on cropping rotations within the following season (Joshi et al., 2007; Mondal et al., 2013). In addition, CIMMYT breeders aim to maintain a range of maturity within the germplasm to allow the development of elite varieties for different growing regions. Therefore, it is important to consider approaches that account for the influence of phenology on grain yield when selecting on VIs.

Selection indices represent a practical breeding strategy to account for these relationships between traits. Selection indices are linear combinations of traits that allow for selection on multiple traits simultaneously on the basis of a single value (reviewed by Céron-Rojas & Crossa, 2018). They are derived using the genetic and phenotypic correlations between traits in conjunction with weights assigned to each trait according to its importance. Restricted selection indices are a type of selection index that enables the improvement of one or more traits while holding other traits constant by setting their weights to zero (Kempthorn & Nordskog, 1959; Openshaw & Hadley, 1984). Restricted selection indices could therefore be used with HTP and phenology records to select indirectly for grain yield while avoiding driving changes in the distribution of phenology.

The primary objective of this study was to assess the potential for utilizing VIs collected with aerial HTP systems at the early-generation seed-limited stage of the breeding program to improve selection for grain yield in wheat.

## MATERIALS AND METHODS

### Field Trial Design

During the 2016-17 and 2017-18 breeding cycles, two sets of 1,008 entries from CIMMYT’s first-year yield trials (YT) were evaluated for agronomic and HTP traits at CENEB in Ciudad Obregón, Mexico. A unique set of 1,008 entries was assessed in each cycle with no overlap in breeding lines between cycles except for check varieties. Each year, the 1,008 entries were arranged in 36 trials containing 28 entries each. Trials were sown in an α-lattice design with 2 replicates and 6 incomplete blocks per replicate, with each block containing 5 entries. Check varieties “Borlaug100 F2014” and “Kachu #1” were included twice per replicate.

Each set of 1,008 entries contained full-sib families ranging in size from 1 to 34 full-sibs with an average of 4 full-sibs per family. In the first replicate of the YT, entries were ordered by pedigree, such that full-sibs were sown adjacent to one another in families in a serpentine order with the check varieties appearing as the first two plots of the replicate. In the second replicate, entries and checks were completely randomized. Plots in the YT consisted of two 80 cm beds containing three rows each for a total plot size of 2.8m × 1.6m.

To empirically evaluate the correlation between small plot performance, including traits measured on these plots, and replicated yield trials, the same 1,008 entries in each breeding cycle were sown in parallel at CENEB as unreplicated 1m × 0.8m small plots (SP) containing three rows each and were evaluated for both agronomic and HTP traits. The 1,008 entries were sown in a serpentine pattern with the order reflecting that of the first replicate of each trial in the YT. During the 2016-17 breeding cycle, the two checks “Borlaug100 F2014” and “Kachu #1” appeared at the beginning of each set of 28 entries. In the 2017-18 breeding cycle, the checks were randomly dispersed within each set of 28 entries. Additional check varieties “MISR 1” and “Baj #1” were added during the 2017-18 breeding cycle and sown ten times each at random within the field.

The YT were sown in pre-irrigated fields on 29 November 2016 and 21 November 2017. The SP were sown in dry soil and received the first irrigation on 25 November 2016 and 23 November 2017. Plots were visually evaluated during the growing season for days to heading (DTHD) and days to maturity (DTMT). DTHD and DTMT were evaluated as the number of days to reach the heading or maturity stages starting from the date of sowing for the YT and from the date of first irrigation for the SP. In the YT, DTHD and DTMT were evaluated on the first replicate only, due to the moderate to high heritability of these traits. Although the breeding program does not typically assess grain yield at the SP stage, both the SP and YT were harvested using plot combine harvesters and then weighed manually for grain yield (GY).

In addition to this proof-of-concept experiment, the SP stage of the formal CIMMYT breeding program was assessed for HTP traits during the 2016-17 breeding cycle. These entries were sown as unreplicated 0.7m × 0.8m beds containing two rows each. Plots were arranged in a serpentine pattern with full-sibs sown adjacent to one another in families. A check variety was sown after every 50 plots, alternating between varieties “Borlaug100 F2014” and “Kachu #1”. Those lines that were visually selected by breeders were promoted to first-year YT in 2017-18. Therefore, the breeding lines evaluated as YT and SP in 2017-18 as part of the proof-of-concept experiment had also been evaluated for HTP as part of the formal SP stage of the breeding program during the previous cycle. These plots are herein referred to as SP_BP_. The SP_BP_ were visually assessed for DTHD and DTMT on a qualitative scale with respect to the nearest sown “Borlaug100 F2014” check. Plots were scored as “early”, “mid”, “late”, and “very late” for DTHD and DTMT. The check “Borlaug 100 F2014” was considered to have medium relative maturity with a score of “mid” for DTHD and DTMT. The first 300 plots of the SP_BP_ were harvested and weighed manually for GY, enabling a direct comparison of the response to selection for visual versus HTP selection.

### Aerial High-Throughput Phenotyping

#### Ground Control Points

The trials were prepared for aerial imaging by distributing ground control points (GCPs) within the field. The GCPs were 27.9 cm × 15.2 cm white polyvinyl sheets mounted on posts. Post heights were 1.2 m in the YT and 1.0 m in the SP. The GPS locations of each GCP were measured with a R4 RTK GPS (Trimble, Sunnyvale, CA) with a horizontal accuracy of 0.025m and a vertical accuracy of 0.035m.

#### Aerial HTP Equipment

A Matrice 100 quadcopter unmanned aerial vehicle (UAV) (DJI, Shenzhen, China) equipped with a MicaSense RedEdge camera (MicaSense, Seattle, WA) was flown over both the YT and SP at multiple time-points during the grain filling developmental growth stage. The Matrice 100 has a maximum takeoff weight of 3600g and can hover for 22-28min with no payload. Following a modification to accommodate an additional battery pack, the maximum no payload hover time was increased to 33-40min. The MicaSense RedEdge weighs 150g and captures one image in 12-bit RAW format per second. Reflectance was recorded at five broadband wavelengths: blue (460-510nm), green (545-575nm), red (630-690nm), near infrared (820-860nm), and red edge (712-722nm). During the 2017-18 breeding cycle, a MicaSense Downwelling Light Sensor (MicaSense, Seattle, WA) integrated with the RedEdge was added on the top of the UAV to improve calibrations when ambient light conditions change throughout the flight.

#### Calibration Panel

Prior to each flight, images of a MicaSense Calibrated Reflectance Panel (MicaSense, Seattle, WA) were captured with the MicaSense RedEdge for use in post-processing of the imagery to account for the ambient light conditions at the time of flight. As provided by MicaSense, the absolute reflectance values of the panel were: 0.366 (blue), 0.366 (green), 0.391 (red), 0.411 (near infrared), and 0.399 (red edge).

#### Acquisition of Aerial Imagery

During the 2016-17 breeding cycle, the UAV was flown with a speed of 12 km/hr at an altitude of 30m, giving an image resolution of 2 cm/pixel. In the 2017-18 breeding cycle, the altitude and speed of the UAV were reduced to 16m and 7 km/hr, increasing the image resolution to 1 cm/pixel. Flights were conducted within 1h of solar noon to minimize variation due to the solar zenith angle (Gu et al., 1992).

#### Aerial Imagery Post-Processing and Data Extraction

A semi-automated data analysis pipeline presented in Haghighattalab et al., (2016) was utilized to analyze each image set and extract plot-level phenotypic values. In summary, the following steps were performed to process each HTP data collection time-point: 1) align the aerial images and building sparse point clouds, 2) import GCP GPS coordinates and geo-reference the images, 3) construct a dense point cloud, 4) create a digital elevation model (DEM), 5) generate an orthomosaic using the DEM, 6) calculate vegetation indices (VIs), and 6) extract the plot-level VI data. The VIs measured were: normalized difference vegetation index (NDVI), green NDVI (GNDVI), and red-edge NDVI (RENDVI). Each VI was calculated following:

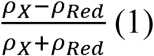

where *ρ*_*Red*_ represents the spectral reflectance recorded in the red region of the light spectrum and *ρ*_*X*_ represents the spectral reflectance recorded in the near-infrared, green, and red-edge regions for NDVI, GNDVI, and RENDVI, respectively.

### Genotyping and Relationship Matrix Development

The 2,016 entries were genotyped with genotyping-by-sequencing (GBS) using PstI-MspI (Poland et al., 2012). The Tassel 5 GBS v2 pipeline was used to process raw Illumina reads and call single nucleotide polymorphism (SNP) markers with minor allele frequency (MAF) of 0.01 (Glaubitz et al., 2014). The unique tags from the Tassel 5 GBS v2 pipeline were aligned to the International Wheat Genome Sequencing Consortium’s RefSeq v1.0 assembly of Chinese Spring (IWGSC, 2018) using Bowtie2 (Langmead & Salzberg 2012) to get a maximum number of tags with unique mapping. The tags with a mapping quality of at least 20 were selected for SNP calling. The SNPs called from the production step were filtered with three criteria: inbreeding coefficient of at least 0.8, Fisher Exact Test (*p-*value<0.001) to determine biallelic single locus SNPs (Poland et al., 2012), and Chi-square test for biallelic segregation with 96 percent expected inbreeding. SNPs that passed at least one filtering criteria were retained, and the subsequent SNP set was filtered to remove those with greater than 50 percent missing data and less than 0.01 MAF. Entries with over 50 percent missing data were also removed from the analysis. The final dataset consisted of 13,271 SNP markers for 1,817 entries. Missing markers were imputed with the marker mean. The genomic relationship matrix (**G**) between entries was calculated according to Endelman and Jannink (2012). The additive pedigree matrix (**A**) was calculated as twice the coefficient of parentage.

### Genetic Value Estimation

Genetic values of each trait for each entry were estimated for the YT and SP experiments separately within each breeding cycle, as well as for the SP_BP_ plots. For the YT, best linear unbiased predictors (BLUPs) were calculated for the agronomic traits and for each VI at each time-point by fitting the following mixed model:

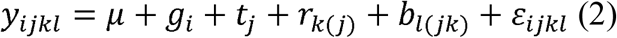

where *y*_*ijkl*_ is the trait value; μ is the overall mean; *g*_*i*_ is the random genetic effect for genotype *i* (BLUP), which are assumed to be independently and identically distributed according to a normal distribution *g*_*i*_∼iid *N*(0, σ_*g*_^2^); *t*_*j*_∼iid *N*(0, σ_*t*_^2^) is the random effect for trial *j*; *r*_*k(j)*_ ∼iid *N*(0, σ_*r*_^2^) is the random effect for replicate *k* within trial *j*; *b*_*l(jk)*_∼iid *N*(0, σ_*b*_^2^) is the random effect for block *l* within replicate *k* and trial *j*; and ε_*ijkl*_ ∼iid *N*(0, σ_*ε*_^2^) is the residual effect. For DTHD and DTMT, which were evaluated on the first replicate only, the random effect for replicate was removed from model (2).

For the SP and SP_BP_ experiments, BLUPs were calculated for the agronomic traits and VIs as by fitting the model:

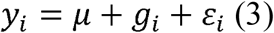

where *y*_*i*_ is the trait value; μ is the overall mean; *g*_*i*_∼iid *N*(0, σ_*g*_^2^) is the random genetic effect for genotype *i*; and *ε*_*i*_ ∼iid *N*(0, σ_ε_^2^***R***_*i*_) is the residual variance for genotype *i*, where ***R***_***i***_ is the correlation matrix for residual effects parameterized as a one-dimensional autoregressive (AR1) process in the column direction to account for potential spatial correlation of observations. An AR1 process model was not applied in the direction of rows as full-sibs were sown adjacent to one another in rows, resulting in confounding between genetic and spatial variation in the row direction.

To account for the effect of phenology, BLUPs for GY and the VIs in the YT and SP were calculated a second time by adding a fixed effect covariate for DTHD to models (2) and (3). BLUPs for the VIs in the SP_BP_ were corrected for DTHD including the qualitative DTHD scores as a fixed effect covariate in model (3).

Significant outliers (*p*-value < 0.05) were identified using Studentized Residuals and were removed from the analysis. In addition, the dataset was subset to remove two families that contained disproportionately high numbers of full-sibs. After further removing entries based on missing genotypic data, the final dataset included 839 and 920 entries in the 2016-17 and 2017-18 breeding cycles, respectively.

To avoid shrinkage on the same data twice (once during the calculation of iid BLUPs and again in the GS models), BLUPs were de-regressed by dividing by their reliability 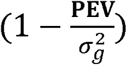, where **PEV** is the prediction error variance of the BLUP and σ_*g*_^2^ is the genotypic variance (Garrick et al., 2009).

Weights for the error variances of the BLUPs were calculated as follows:

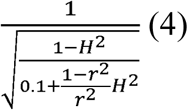

where *H*^*2*^ is the proportion of the total variance explained by the genotypic variance component, σ_*g*_^2^, and *r*^*2*^ are the reliabilities of the BLUPs (Garrick et al., 2009). These weights were used in downstream genomic prediction analyses to preserve information about heterogeneous variances driven by the differences in plot size and replication between the YT and SP. Without weights, this information would be otherwise ignored in a two-step genomic prediction procedure.

### Heritability, Trait Correlation, and Response to Selection

To estimate the narrow-sense heritability, variance components for each of the agronomic and HTP traits were estimated according to the following model:

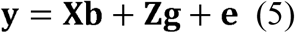

where **y** is a vector of de-regressed BLUPs calculated from models (2) or (3); **X** is the fixed effects design matrix; **b** is a vector of fixed effects; **Z** is the random effects design matrix; **g** is a vector of random effects for genotype estimated assuming **g**∼*N*(0, **G**σ_*a*_^2^) where **G** is the genomic relationship matrix and σ_*a*_^2^ is the additive genetic variance; and **e** is a vector of residuals where **e**∼iid *N*(0, **I**σ_*e*_^2^), σ_*e*_^2^ is the residual variance, and **I** is the identity matrix. Narrow-sense heritability was calculated as 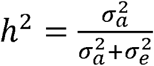.

Trait phenotypic correlations were calculated as the Pearson’s correlations between the iid BLUPs for each trait derived from models (2) and (3). Variance components for deriving the genetic correlations between each pair of traits were calculated according to the bivariate model:

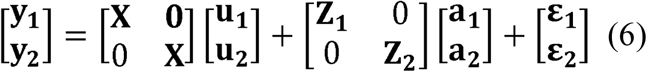

where **y**_**1**_ and **y**_**2**_ are vectors of de-regressed BLUPs for each trait calculated from models (1) or (2); **X** is the fixed effects design matrix, which is the same for both traits; **u**_**1**_ and **u**_**2**_ are vectors of fixed effects for each trait; **Z**_**1**_ and **Z**_**2**_ are the random effects design matrices for each trait; **a**_**1**_ and **a**_**2**_ are vectors of random effects for each trait; and **ε**_**1**_ and **ε**_**2**,_ are vectors of residuals for each trait. The model was fit assuming 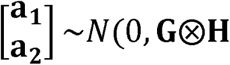, where **G** is the genomic relationship matrix and **H** is the variance-covariance matrix for the two traits. For the residual, 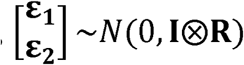, where **I** is the identity matrix and **R** is the residual variance-covariance matrix for the two traits. The resulting variance components were used to calculate the genetic correlation between each pair of traits according to:

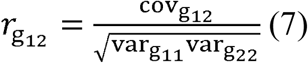

where 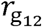 and 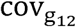 are the genetic correlation and covariance, respectively, between the traits, and 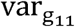 and 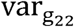 are the genetic variances for each trait.

Response to selection was assessed by identifying the top performing breeding lines according to traits measured in the SP and calculating the percent difference in YT trait values between the means of those superior breeding lines and the total population. Response to selection was estimated for selection intensities from 100 (no selection) to 20 percent of breeding lines. Response to visual and HTP selection in the SP_BP_ was measured by calculating the percent difference in mean GY of the SP_BP_ between the selected lines and the total population.

### Selection Schemes

A train-test (TRN-TST) partitioning scheme was developed to estimate model accuracy within each breeding cycle. In each partition, 80 percent of the total number of lines within a breeding cycle was sampled to form the TRN set, while the remaining 20 percent of entries formed the TST set. Whole families were grouped during sampling to avoid full-sibs from the same family appearing in both the TRN and TST sets, which greatly inflates this prediction accuracy but does not represent a real-world implementation for forward selection into new breeding crosses.

#### High-Throughput Phenotyping Selection

To assess the ability to predict GY of the YT using solely HTP information from the SP stage, the following linear regression model was fit using GY and HTP information from the YT for the entries in the TRN set:

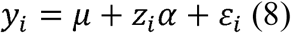

where *y*_*i*_ is the de-regressed BLUP of genotype *i* for GY in the YT calculated from model (2), μ is the overall mean, *z*_*i*_ is the de-regressed BLUP of genotype *i* for an VI time-point observed in the YT calculated from model (2), *α* is the regression coefficient for the HTP trait, and *ε*_*i*_ is the residual for genotype *i*.

This model was then applied to predict GY for the breeding lines in the TST set. The de-regressed BLUPs for the same VI time-point observed in the SP calculated from model (3) were assigned to *z*_*i*_ while values for GY were “hidden.” Models were assessed for each VI on each HTP time-point.

#### Univariate Genomic Selection

As a basis for comparison, univariate GS models using genomic marker or pedigree information were developed to predict GY for each breeding cycle following the form:

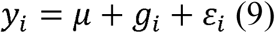

where *y*_*i*_ are the de-regressed BLUPs for GY in the YT calculated in model (2); μ is the overall mean; *g*_*i*_ is the random effect for genotype *i* with *g*_*i*_∼*N*(0, **G**σ_*g*_^2^) or *g*_*i*_∼*N*(0, **A**σ_*g*_^2^), where **G** is the genomic relationship matrix and **A** is the pedigree matrix; and *ε*_*i*_ ∼iid *N*(0, **R**σ_*ε*_^2^) is the residual error for genotype *i* where **R** is the residual variance matrix. Weights from model (4) were included in the diagonal of **R**. These models contained no information on HTP traits.

#### Multi-Trait Genomic Selection

Finally, multi-trait GS approaches integrating HTP records with genomic/pedigree information to predict GY were developed following the bivariate model (6) where **y**_**1**_ is a vector of de-regressed BLUPs for GY in the YT calculated from model (2); **y**_**2**_ is a vector of de-regressed BLUPs for a VI time-point observed in the YT calculated from model (2); **X** is the fixed effects design matrix, which is the same for each trait; **u**_**1**_ and **u**_**2**_ are vectors of fixed effects for GY and the HTP time-point; **Z**_**1**_ and **Z**_**2**_ are the random effects design matrices for GY and the VI time-point; **a**_**1**_ and **a**_**2**_ are vectors of random effects for GY and the VI time-point; and **ε**_**1**_ and **ε**_**2**_ are vectors of residuals for GY and the VI trait. This model was fit assuming 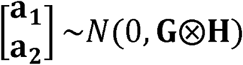 or 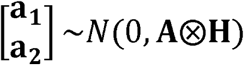, where **G** is the genomic relationship matrix, **A** is the pedigree matrix, and **H** is the variance-covariance matrix for GY and the VI time-point. The weights from model (4) were included in the diagonal of **R**.

This model was then applied to the breeding lines in the TST set, for which the de-regressed BLUPs for the same HTP time-point observed in the SP calculated from model (3) represented **y**_**2**_ while the values for GY were masked. All possible combinations of HTP time-points, VIs, and relationship matrices were tested.

The same 20 TRN-TST partitions were evaluated for HTP selection, univariate GS, and multi-trait GS to ensure fair comparisons between methods. Predictive abilities were calculated by taking the Pearson’s correlation of the predicted values for the TST set lines with the iid BLUPs for GY in the YT from model (2). The reported predictive abilities are represented by the mean and standard deviation of the 20 TRN-TST partitions.

### Accounting for Phenology

Due to the strong associations between GY, VIs, and phenology observed in this population, two approaches were evaluated to account for the influence of phenology on GY. In the first approach, iid BLUPs for DTHD observed in the YT calculated from model (2) were included in prediction models (8), (9), and (5) as a fixed effect covariate. For model validation, the predicted values were correlated to iid BLUPs for GY in the YT that had likewise been corrected for DTHD by including DTHD records observed in the YT as a fixed effect covariate in model (2). Prediction models (8), (9), and (5) were tested both with and without this correction for DTHD.

In the second approach, a desired gains selection index taking into account the genetic correlation between GY and DTHD was developed to assess the ability of prediction models (8), (9), and (5) to identify breeding lines with high GY without selecting directionally on DTHD. The desired gains index takes the form:

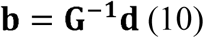

where **b** is a vector of the selection index weights, **G**^**−1**^ is the inverse of the genetic variance-covariance matrix among traits, and □ is a vector of the desired gains for the traits (Pešek & Baker, 1969).

To avoid directional selection on DTHD, the desired gain for DTHD, *d*_*DTHD*_, was assigned to zero. In each breeding cycle, the desired gain for GY, *d*_*GY*_, was set equal to one standard deviation of the iid BLUPs for GY in the YT. **G**^**−1**^ was calculated by fitting the bivariate model (5) with the genetic relationship matrix **G** and deregressed BLUPs for GY and DTHD in the YT. **G**^**−1**^ was calculated separately for each breeding cycle, using records for all lines. The index weights, **b**, were then derived in model (10) using the resulting values for **G**^**−1**^ and assigned values for **d**.

To validate the prediction models in the context of the restricted selection index, models (8), (9), and (5) were fit as described to predict GY for the TST set. The GY predictions were then combined with observed records for DTHD in the SP to obtain the “predicted” restricted gains index values:

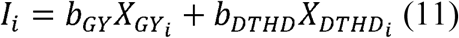

where *I*_*i*_ is the index value for genotype *i*; *b*_*GY*_ and *b*_*DTHD*_ are the index weights for GY and DTHD, respectively, derived from model (10); 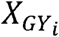 is the predicted value for GY of genotype *i* from prediction models (8), (9), and (5); and 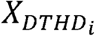 is the observed iid BLUP for DTHD in the SP for genotype *i* (Smith, 1936; Hazel, 1943).

For model validation, the observed index values for all breeding lines were calculated with model (11) using the observed iid BLUPs for GY and DTHD in the YT derived from model (2). Predictive abilities are represented as the Pearson’s correlations between the predicted and observed index values for the lines in the TST set.

### Software

Image processing was conducted using Python, open source software QGIS (QGIS Development Team, 2019, QGIS Geographic Information System, Open Source Geospatial Foundation Project, http://qgis.osgeo.org), and Agisoft PhotoScan (Agisoft LLC, St. Petersburg, Russia). Data analyses were implemented in the R environment (R Core Team, 2016, Vienna, Austria) with the package “ASReml-R” (Gilmour et al., 2014) for R. The genomic relationship matrix (**G**) was calculated using the A.mat() function within the R package “rrBLUP” (Endelman, 2011). The coefficients of parentage for calculating the pedigree relationship matrix (**A**) were estimated with the “Browse” application of the International Crop Information System software (McLaren et al., 2000).

## Data Availability

All phenotypic and genotypic data required to confirm the results presented in this study are available on CIMMYT Dataverse (http://hdl.handle.net/11529/10548379).

## RESULTS

The distributions of BLUPs for GY in the YT were consistent across the two breeding cycles, while BLUPs for GY in the SP were more variable within and across cycles (Table 1). Both the YT and SP reached the heading and maturity stages around the same time during both breeding cycles (Fig. 1) and were therefore approximately at the same stage of development when the VIs were recorded. The YT and SP were phenotyped for VIs with the UAV at two time-points (TP-1: 13 Mar; TP-2: 20 Mar) in 2016-17 and at three time-points (TP-1: 07 Mar; TP-2: 12 Mar in the SP, 13 Mar in the YT; TP-3: 19 Mar) in 2017-18 (Fig. 1). The SP_BP_ was phenotyped with the UAV on the same time-points as the YT and SP in 2016-17.

**Table 1.** Trait correlations within and between lines in the YT, SP, and SP_BP_

| 2016-17 |  | YT |  |  |  |  | SP |  |  |  |  |
| --- | --- | --- | --- | --- | --- | --- | --- | --- | --- | --- | --- |
|  |  | GY | DTHD | DTMT | NDVI TP-1 | NDVI TP-2 | GY | DTHD | DTMT | NDVI TP-1 | NDVI TP-2 |
| YT | GY | 6.45<br>(0.32) | 0.41<br>(0.61) | 0.44<br>(0.62) | 0.50<br>(0.72) | 0.51<br>(0.72) | 0.32<br>(0.67) | 0.40<br>(0.63) | 0.42<br>(0.69) | 0.43<br>(0.72) | 0.44<br>(0.73) |
|  | DTHD | - | 74.5<br>(4.2) | 0.78<br>(0.93) | 0.74<br>(0.82) | 0.80<br>(0.89) | 0.08<br>(0.12) | 0.82<br>(0.96) | 0.65<br>(0.85) | 0.60<br>(0.62) | 0.71<br>(0.81) |
|  | DTMT | - | - | 122.5<br>(3.1) | 0.72<br>(0.80) | 0.80<br>(0.90) | 0.12<br>(0.17) | 0.73<br>(0.89) | 0.69<br>(0.91) | 0.59<br>(0.67) | 0.72<br>(0.86) |
|  | NDVI TP-1 | 0.32<br>(0.48) | - | - | 0.824<br>(0.021) | 0.94<br>(0.95) | 0.17<br>(0.19) | 0.72<br>(0.75) | 0.69<br>(0.79) | 0.75<br>(0.89) | 0.78<br>(0.86) |
|  | NDVI TP-2 | 0.34 (0.45) | - | - | 0.85<br>(0.79) | 0.669<br>(0.053) | 0.14<br>(0.05) | 0.78<br>(0.82) | 0.76<br>(0.86) | 0.76<br>(0.88) | 0.84<br>(0.92) |
| SP | GY | 0.30 (0.78) | - | - | 0.14<br>(0.41) | 0.09<br>(0.21) | 5.20<br>(0.64) | 0.12<br>(0.05) | 0.14<br>(0.12) | 0.25<br>(0.27) | 0.20<br>(0.16) |
|  | DTHD | - | - | - | - | - | - | 78.2<br>(3.5) | 0.76<br>(0.85) | 0.70<br>(0.64) | 0.81<br>(0.81) |
|  | DTMT | - | - | - | - | - | - | - | 121.3<br>(3.5) | 0.75<br>(0.84) | 0.86<br>(0.95) |
|  | NDVI TP-1 | 0.22 (0.54) | - | - | 0.48<br>(0.89) | 0.44<br>(0.91) | 0.23<br>(0.33) | - | - | 0.774<br>(0.038) | 0.91<br>(0.93) |
|  | NDVI TP-2 | 0.17 (0.47) | - | - | 0.42<br>(0.74) | 0.49<br>(0.89) | 0.17<br>(0.21) | - | - | 0.82<br>(0.90) | 0.569<br>(0.082) |

| 2017-18 |  | YT |  |  |  |  |  | SP |  |  |  |  |  | SP<sub>BP</sub> |  |
| --- | --- | --- | --- | --- | --- | --- | --- | --- | --- | --- | --- | --- | --- | --- | --- |
|  |  | GY | DTHD | DTMT | NDVI TP-1 | NDVI TP-2 | NDVI TP-3 | GY | DTHD | DTMT | NDVI TP-1 | NDVI TP-2 | NDVI TP-3 | NDVI TP-1 | NDVI TP-2 |
| YT | GY | 6.24<br>(0.32) | 0.43<br>(0.60) | 0.45<br>(0.64) | 0.46<br>(0.64) | 0.45<br>(0.60) | 0.49<br>(0.63) | 0.40<br>(0.89) | 0.42<br>(0.57) | 0.39<br>(0.58) | 0.37<br>(0.55) | 0.39<br>(0.60) | 0.42<br>(0.57) | 0.33<br>(0.53) | 0.36<br>(0.52) |
|  | DTHD | - | 79.0<br>(3.6) | 0.84<br>(0.90) | 0.73<br>(0.86) | 0.79<br>(0.88) | 0.80<br>(0.89) | 0.19<br>(0.52) | 0.88<br>(0.96) | 0.66<br>(0.85) | 0.67<br>(0.76) | 0.73<br>(0.83) | 0.80<br>(0.88) | 0.51<br>(0.65) | 0.61<br>(0.73) |
|  | DTMT | - | - | 126.1<br>(2.6) | 0.69<br>(0.83) | 0.74<br>(0.85) | 0.79<br>(0.89) | 0.16<br>(0.46) | 0.76<br>(0.85) | 0.68<br>(0.89) | 0.63<br>(0.71) | 0.66<br>(0.79) | 0.74<br>(0.84) | 0.51<br>(0.67) | 0.63<br>(0.77) |
|  | NDVI TP-1 | 0.23<br>(0.31) | - | - | 0.491<br>(0.022) | 0.88<br>(0.95) | 0.87<br>(0.93) | 0.25<br>(0.65) | 0.67<br>(0.82) | 0.57<br>(0.81) | 0.72<br>(0.90) | 0.73<br>(0.92) | 0.71<br>(0.88) | 0.51<br>(0.72) | 0.56<br>(0.73) |
|  | NDVI TP-2 | 0.19<br>(0.21) | - | - | 0.72<br>(0.83) | 0.504<br>(0.023) | 0.89<br>(0.94) | 0.23<br>(0.53) | 0.72<br>(0.85) | 0.62<br>(0.83) | 0.73<br>(0.87) | 0.77<br>(0.92) | 0.78<br>(0.92) | 0.62<br>(0.78) | 0.65<br>(0.79) |
|  | NDVI TP-3 | 0.24<br>(0.28) | - | - | 0.69<br>(0.70) | 0.71<br>(0.74) | 0.355<br>(0.035) | 0.24<br>(0.66) | 0.75<br>(0.86) | 0.67<br>(0.89) | 0.69<br>(0.81) | 0.74<br>(0.88) | 0.79<br>(0.92) | 0.56<br>(0.73) | 0.64<br>(0.81) |
| SP | GY | 0.35<br>(0.84) | - | - | 0.15<br>(0.43) | 0.13<br>(0.22) | 0.13<br>(0.52) | 7.69<br>(0.45) | 0.17<br>(0.60) | 0.17<br>(0.57) | 0.20<br>(0.58) | 0.23<br>(0.60) | 0.23<br>(0.88) | 0.22<br>(0.46) | 0.23<br>(0.44) |
|  | DTHD | - | - | - | - | - | - | - | 77.1<br>(3.8) | 0.73<br>(0.91) | 0.70<br>(0.79) | 0.76<br>(0.84) | 0.85<br>(0.60) | 0.49<br>(0.67) | 0.60<br>(0.74) |
|  | DTMT | - | - | - | - | - | - | - | - | 125.8<br>(2.4) | 0.65<br>(0.83) | 0.71<br>(0.89) | 0.81<br>(0.84) | 0.53<br>(0.80) | 0.63<br>(0.88) |
|  | NDVI TP-1 | 0.09<br>(0.24) | - | - | 0.46<br>(0.80) | 0.46<br>(0.73) | 0.32<br>(0.51) | 0.11<br>(0.26) | - | - | 0.534<br>(0.021) | 0.89<br>(0.95) | 0.82<br>(0.89) | 0.53<br>(0.72) | 0.56<br>(0.70) |
|  | NDVI TP-2 | 0.09<br>(0.28) | - | - | 0.42<br>(0.78) | 0.48<br>(0.82) | 0.34<br>(0.61) | 0.15<br>(0.26) | - | - | 0.78<br>(0.90) | 0.491<br>(0.027) | 0.89<br>(0.95) | 0.58<br>(0.78) | 0.62<br>(0.79) |
|  | NDVI TP-3 | 0.09<br>(0.19) | - | - | 0.28<br>(0.60) | 0.42<br>(0.76) | 0.40<br>(0.76) | 0.15<br>(0.18) | - | - | 0.59<br>(0.61) | 0.73<br>(0.82) | 0.397<br>(0.047) | 0.62<br>(0.73) | 0.71<br>(0.86) |
| SP<sub>BP</sub> | NDVI TP-1 | 0.11<br>(0.26) | - | - | 0.17<br>(0.33) | 0.34<br>(0.50) | 0.20<br>(0.31) | 0.12<br>(0.27) | - | - | 0.21<br>(0.34) | 0.28<br>(0.52) | 0.34<br>(0.54) | 0.759<br>(0.035) | 0.85<br>(0.95) |
|  | NDVI TP-2 | 0.09<br>(0.20) | - | - | 0.13<br>(0.22) | 0.27<br>(0.39) | 0.22<br>(0.37) | 0.11<br>(0.21) | - | - | 0.14<br>(0.26) | 0.21<br>(0.43) | 0.34<br>(0.62) | 0.77<br>(0.90) | 0.549<br>(0.068) |

YT, yield trials; SP, small plots; SP<sub>BP</sub>, small plots from the 2016-17 breeding program; GY, grain yield; DTHD, days to heading; DTMT, days to maturity; NDVI, normalized difference vegetation index; TP, time-point. The diagonals contain the mean of the iid BLUPs for each trait derived from models (1) and (2). The values in parentheses along the diagonal are the standard deviations of the iid BLUPs for each trait. The upper triangles contain the Pearson's correlation between each pair of traits. The lower triangles contain the Pearson's correlations among and between GY and the NDVI time-points following correction for DTHD. The values in parentheses in the upper and lower triangles are the genetic correlations between each pair of traits.

**Figure 1.**
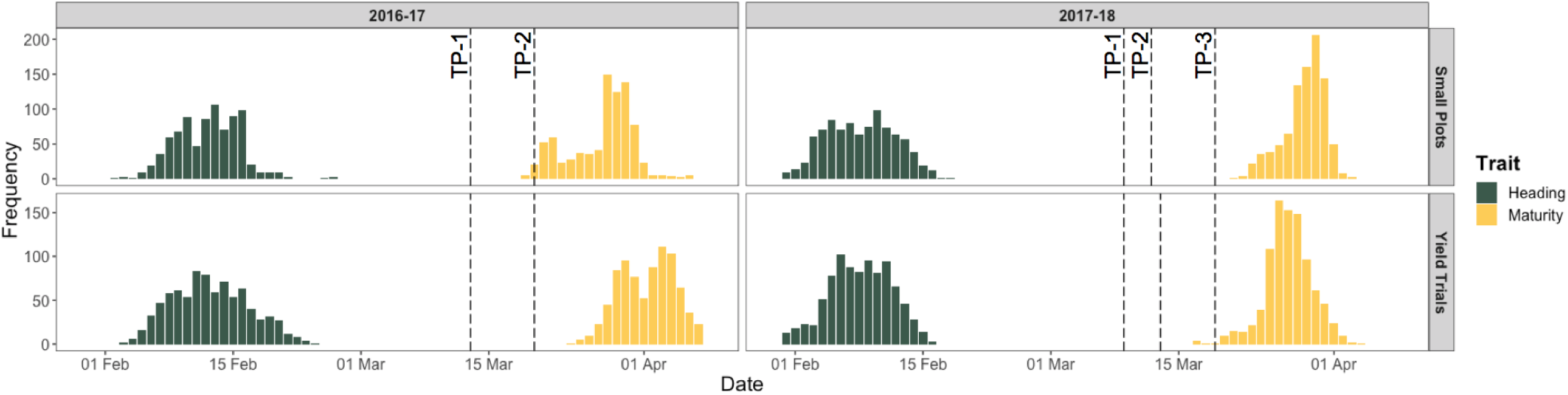
UAV phenotyping time-points in relationship to heading and maturity dates in each experiment. YT, yield trials; SP, small plots; TP, time-point; UAV, unmanned aerial vehicle; VI, vegetation index. The histograms represent the heading and maturity dates of the SP and YT in 2016-17 and 2017-18. The dotted red lines indicate the time-points on which each experiment was phenotyped for VIs with the UAV.

### Trait Heritabilities

As expected, heritability estimates for GY were considerably greater for the YT than for the SP in both the 2016-17 and 2017-18 cycles (Table 2). Accounting for variation in DTHD resulted in a slight decrease in the heritability of GY in the YT and in the SP from 2016-17 but had no effect in the SP from 2017-18. For a given VI time-point, the heritability estimates for the three VIs differed from one another by an average of 0.03. Due to the overall similarity between VIs for heritability, only the estimates for NDVI are reported.

**Table 2.** Narrow-sense heritability (*h*^*2*^) estimates for the agronomic and NDVI traits

| Breeding Cycle | Trait | YT $h^2$ | SP $h^2$ | SP<sub>BP</sub> $h^2$ |
| --- | --- | --- | --- | --- |
| 2016-17 | GY | 0.55 (0.46) | 0.39 (0.40) | - |
|  | DTHD | 0.79 | 0.82 | - |
|  | DTMT | 0.65 | 0.80 | - |
|  | NDVI TP-1 | 0.77 (0.58) | 0.69 (0.75) | 0.79 (0.65) |
|  | NDVI TP-2 | 0.84 (0.57) | 0.79 (0.78) | 0.83 (0.65) |
| 2017-18 | GY | 0.72 (0.65) | 0.37 (0.31) | - |
|  | DTHD | 0.82 | 0.85 | - |
|  | DTMT | 0.83 | 0.65 | - |
|  | NDVI TP-1 | 0.79 (0.59) | 0.82 (0.74) | - |
|  | NDVI TP-2 | 0.79 (0.58) | 0.78 (0.65) | - |
|  | NDVI TP-3 | 0.83 (0.64) | 0.83 (0.59) | - |

YT, yield trials; SP, small plots; SP<sub>BP</sub>, small plots from the 2016-17 breeding program;
GY, grain yield; DTHD, days to heading; DTMT, days to maturity; NDVI, normalized
difference vegetation index; TP, time-point. Values in parentheses are the narrow-sense
heritability for the traits corrected for DTHD.

Heritability estimates for the NDVI time-points were similar across the YT and SP and had an average value of 0.79, which was comparable to the heritabilities of DTHD and DTMT and considerably exceeded heritability estimates for GY. As with GY, accounting for DTHD in the NDVI time-points resulted in a slight reduction in heritability. Despite the smaller plot size in the SP_BP_ compared to the SP, heritability estimates for the NDVI time-points in the SP_BP_ experiment were similar to those for the NDVI time-points recorded in the SP on the same time-points.

### Trait Phenotypic and Genetic Correlations

Strong phenotypic and genetic correlations were observed among and between the agronomic and VI traits within the YT, SP, and SP_BP_ (Table 1). The phenotypic and genetic correlations for each pair of traits largely followed the same trends, and therefore only the phenotypic correlations are discussed herein. The strongest correlations were observed between the three VIs recorded on the same time-point, which indicates that there was minimal difference in the information captured among the VIs. As a result, only correlations using NDVI, the most commonly used VI in the context of plant breeding among those tested, are reported and discussed. Very high correlations were also observed for NDVI recorded on different time-points, which suggests that changes in canopy reflectance during the period in which VIs were measured were marginal.

Despite the small plot size and lack of replication in the SP, the correlation for GY between the SP and YT was relatively moderate, though correlations between NDVI and GY were lower in the SP than in the YT. The correlations between NDVI in the SP and GY in the YT ranged from 0.37 to 0.44 and was only slightly reduced for SP_BP_, which had a smaller plot size than the SP and was measured in a different breeding cycle than GY in the YT.

Correcting for DTHD had a minimal impact on the correlation between NDVI and GY when measured both traits were measured in the SP. The effect was more pronounced for the YT as well as when NDVI and GY were evaluated in the SP and YT, respectively. Strong relationships were also observed between the NDVI time-points and DTHD/DTMT, averaging 0.76 for the YT and SP. Notably, DTHD and DTMT observed in both the YT and SP was moderately predictive of GY in the YT, with correlations ranging from 0.39 to 0.45.

### Selection on High-Throughput Phenotyping Traits

By identifying the top breeding lines in terms of NDVI assessed in the SP and SP_BP_, results show that using NDVI as the selection criterion at the early generation seed-limited stage may result in considerable improvements in GY in the YT (Fig. 2), with projected increases of 2.17, 1.68, and 0.97 percent at selection intensities of 25, 50, and 75 percent, respectively. This approach was observed to more effective than selecting on GY measured in the SP during the 2016-17 cycle, though the two approaches showed similar results for the 2017-18 cycle. However, selecting on NDVI measured in the SP or SP_BP_ would have resulted in significant directional response in DTHD in the YT, primarily favoring later-maturing breeding lines. Conversely, selecting on GY measured in the SP would have had no effect on DTHD in the YT.

**Figure 2.**
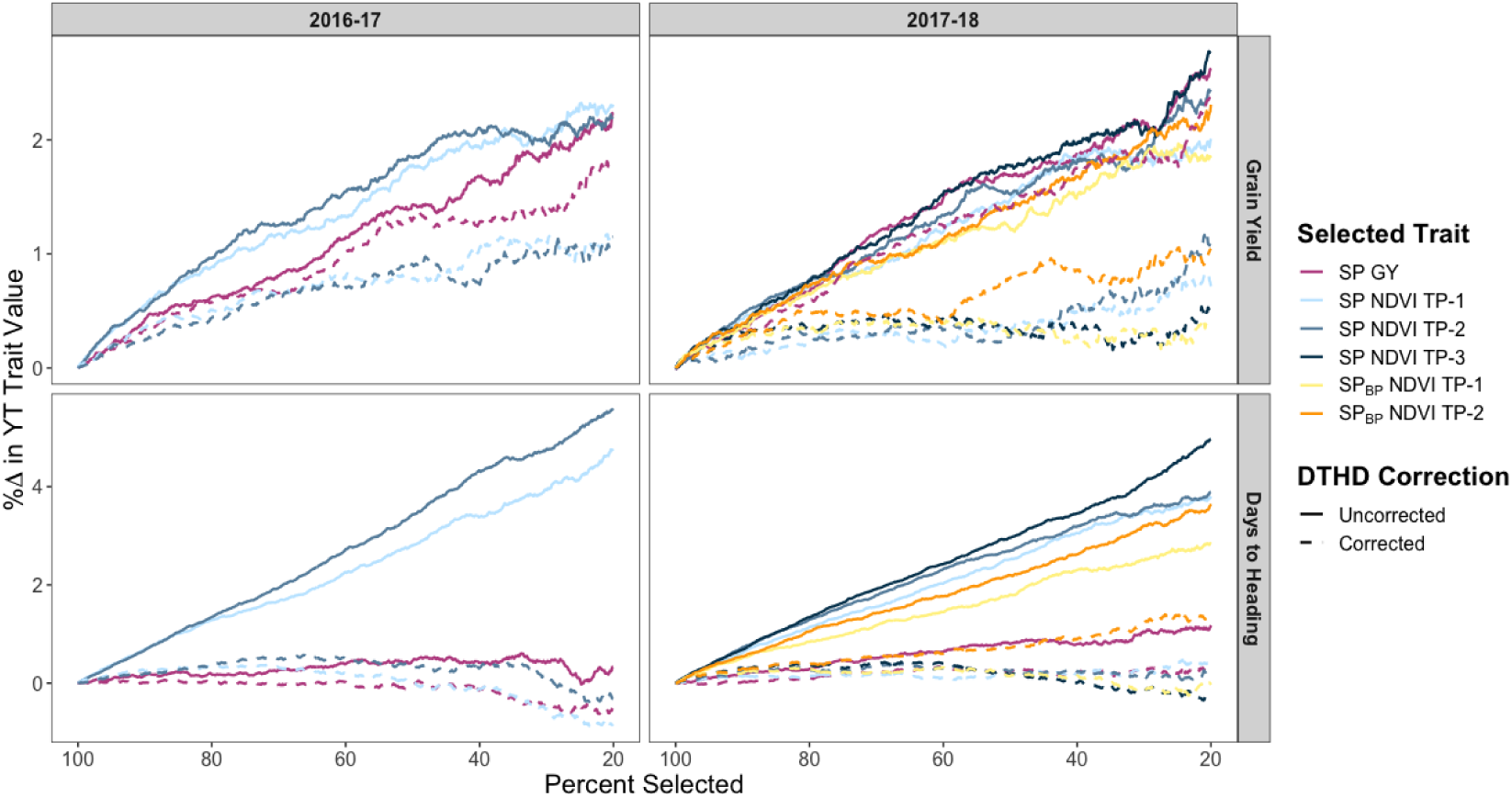
Response to selection in the YT when selecting on traits measured in the SP and SP_BP_. SP_BP_, small plots from the 2016-17 breeding program; GY, grain yield; NDVI, normalized difference vegetation index; DTHD, days to heading. The x-axis represents the selection intensity expressed as the percentage of the population selected based on the SP or SP_BP_ trait. The y-axis represents the corresponding percent change in the YT trait value.

When accounting for DTHD in the NDVI measurements in the SP, the response to selection for GY in the YT was reduced to below 1 percent at most levels of selection intensity. Accounting for DTHD in the GY records in the SP also reduced the projected response to selection for GY in the YT, however the decrease was less pronounced. As expected, selecting on DTHD-corrected NDVI or GY records from the SP would have had minimal impact on the distribution of DTHD in the YT.

### Visual Versus High-Throughput Phenotyping Selection

Prior to harvest in 2016-17, the breeders made visual selections within the first 300 plots of the SP_BP_ for promotion to the first-year YT in the 2017-18 breeding cycle. Of the 293 breeding lines (excluding checks), 36 were selected, representing a selection intensity of 12.3 percent. GY was measured on both the unselected and selected SP_BP_ plots, and results showed that visual selection provided a 0.38 percent response to selection. Among the total population of 293, over half were scored as “early” while the remaining were divided relatively evenly among “mid” and “late” scores for DTHD, with less than 2 percent scored as “very late”. The distribution for DTHD among the selected lines approximately reflected this, though all “very late” lines were eliminated.

In the SP_BP_, the NDVI time-points had correlations with GY of 0.13 for TP-1 and 0.05 for TP-2 (Fig. 3A). Despite these weak relationships, NDVI was effective in identifying SP_BP_ breeding lines with low GY for culling. If the level of selection intensity that the breeders used for visual selection had been applied to the SP_BP_ using the NDVI time-points as the selection criteria, the response to selection in GY of the SP_BP_ would have been 1.54 and 1.40 percent for TP-1 and TP-2, respectively. Of the top 36 lines in terms of NDVI, 25 would have been selected by both TP-1 and TP-2. Only 4 and 5 of the lines visually selected by breeders were among the top 36 in terms of NDVI TP-1 and TP-2, respectively. However, over half of these lines were scored as “late” or “very late” for DTHD.

**Figure 3.**
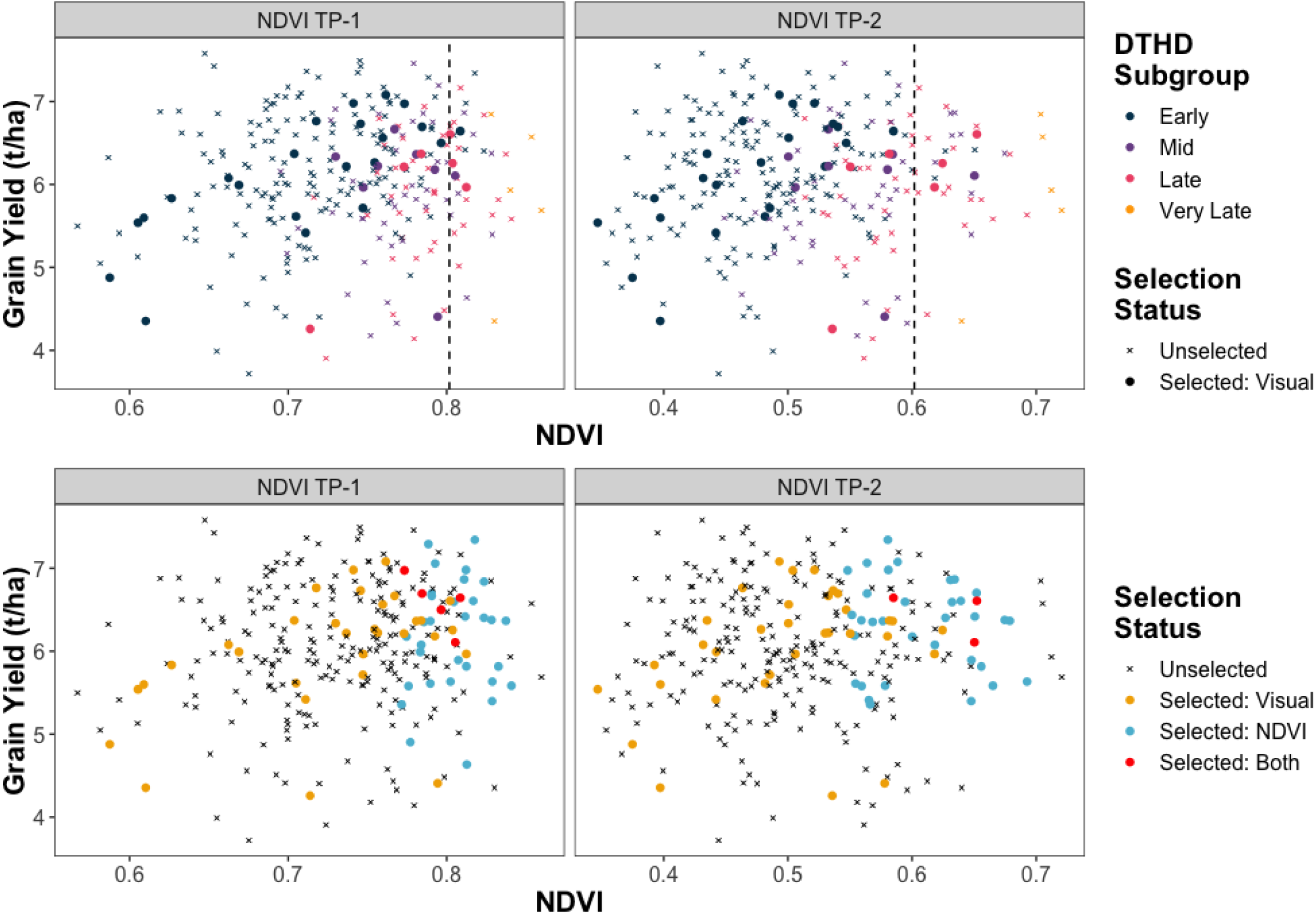
Visual versus HTP selection in the SP_BP_. The x-axis shows NDVI values measured in the SP_BP_. The y-axis represents GY values assessed in the SP_BP_. A. The color represents the qualitative DTHD subgroups the breeding lines were assigned to. The shape indicates which lines were visually selected by breeders. Points to the right of the dashed lines indicate lines that would have been selected using NDVI as the selection criteria, given the same level of selection intensity used by the breeders for visual selection. B. Directional selection on DTHD is avoided by using NDVI to select the top 22, 8, and 6 lines in the “early”, “mid”, and “late” DTHD subgroups, respectively. The color and shape indicate if which lines were selected, and, if so, by which selection method(s).

Since CIMMYT breeders attempt to avoid selecting directionally on DTHD when performing visual selection, a more appropriate comparison would be to identify the top breeding lines in terms of NDVI for each qualitative DTHD score. The breeders visually selected 22, 8, and 6 lines scored as “early,” “mid,” and “late” for DTHD, respectively. By taking subsets of the “early,” “mid,” and “late” lines and identifying the top 22, 8, and 6 lines, respectively, based on NDVI, the response to selection in GY for the SP_BP_ would have been 2.91 percent for TP-1 and 4.23 percent for TP-2 (Fig. 3B). There was still minimal overlap between the lines selected by the breeders and the top lines for NDVI using this approach.

### Predictive Abilities

The predictive abilities of the three VIs gave similar results with no individual VI providing a consistent, significant advantage. In addition, the predictive abilities of models using the pedigree relationship matrix were low, with an average of 0.16 for the univariate GS models, likely due to the large numbers of full-sibs and missing information in the pedigree records. Therefore, only the predictive abilities of models using NDVI and the genomic relationship matrix (**G**) are reported and discussed further. When using NDVI traits alone to predict yield in the absence of genomic marker or pedigree information, predictive abilities were moderate, averaging 0.43 and 0.38 in 2016-17 and 2017-18, respectively, with minimal variation observed between NDVI time-points (Fig. 4). This exceeded predictive abilities for univariate GS, though multi-trait GS models that incorporated both NDVI and genomic information provided the highest predictive abilities overall.

**Figure 4.**
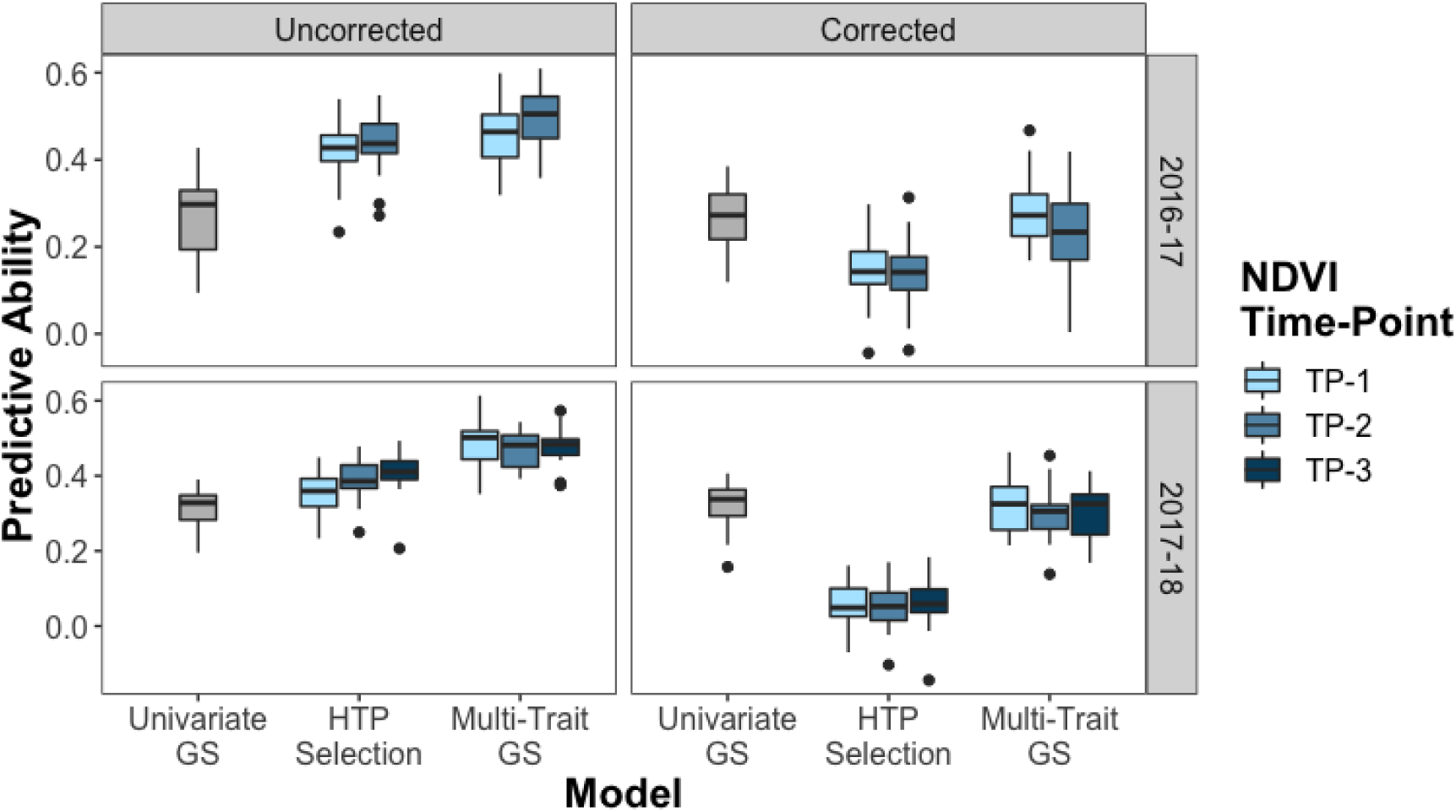
Predictive abilities of univariate GS, HTP selection, and multi-trait GS uncorrected and corrected for phenology. GS, genomic selection; HTP, high-throughput phenotyping. Predictive ability is expressed as the Pearson’s correlations between predictions and observed iid BLUPs for GY in the YT across 20 random TRN-TST iterations. The univariate and multi-trait GS models were fit using the genomic relationship matrix (**G**). Corrected predictive abilities were estimated using genetic values that were calculated with DTHD as a fixed effect covariate.

When estimating model predictive abilities, two approaches were taken to account for confounding of DTHD with both NDVI and GY. In the first approach, iid BLUPs for DTHD in the YT were included as a fixed effect in the prediction model. This correction had a negligible impact on univariate GS, but the predictive abilities of the HTP selection models were strongly reduced to the extent that they became less predictive than univariate GS. Following the correction, the predictive abilities of the multi-trait GS models were more similar to those of univariate GS.

The second approach was to develop a restricted selection index to identify superior breeding lines for GY without selecting directionally on DTHD while taking into account the genetic correlation between the traits (Fig. 5). The observed index values showed phenotypic correlations of 0.89 and 0.85 with GY in 2016-17 and 2017-18, respectively, conferring a response to selection of 0.44 in both breeding cycles. As desired, the index had a negligible relationship with DTHD, with correlations of -0.05 and -0.10 in 2016-17 and 2017-18, respectively. The narrow-sense heritability of the index was 0.47 in 2016-17 and 0.64 in 2017-18. Genetic correlations between the index and the NDVI traits in the SP were moderate during the 2016-17 breeding cycle, with values 0.47 and 0.38 for TP-1 and TP-2, respectively. However, low correlations were observed in 2017-18, with values of 0.13, 0.14, and 0.05 for TP-1, TP-2, and TP-3, respectively.

**Figure 5.**
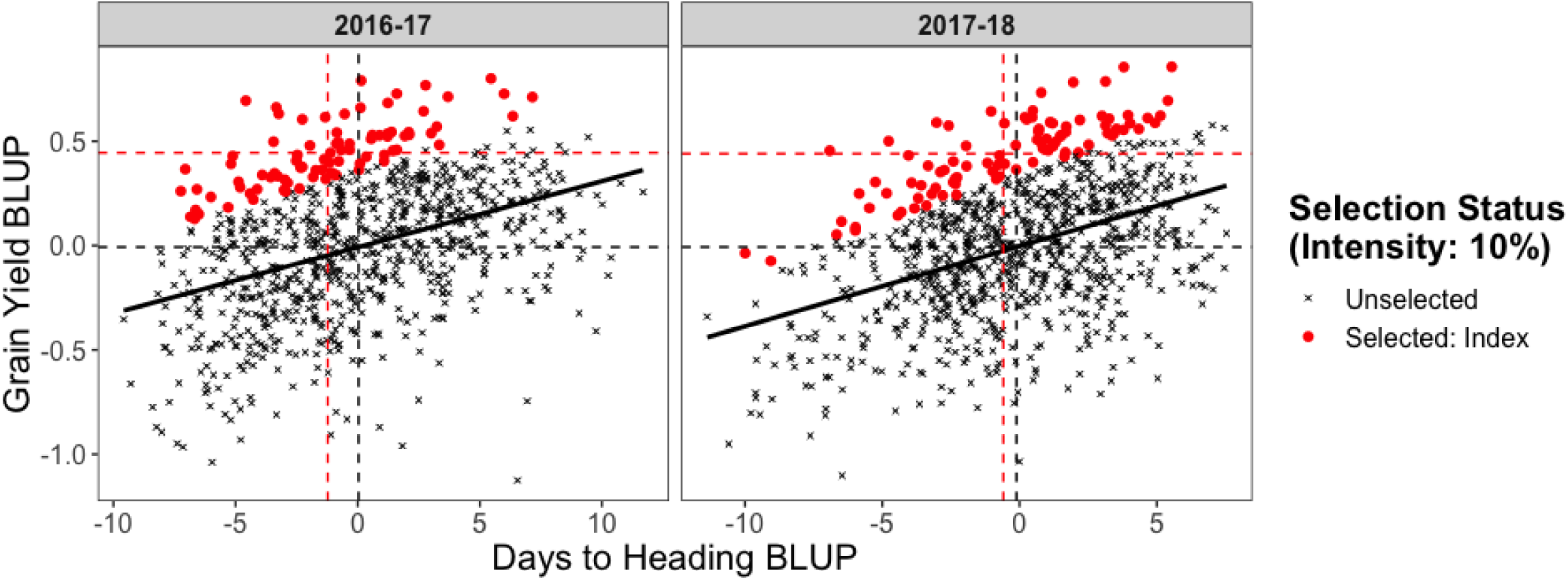
Restricted selection index to increase GY while holding DTHD constant. Plotted points are the iid BLUPs for DTHD and GY in the YT in 2016-17 and 2017-18. The color and shape of the points indicate which breeding lines were selected based on the restricted selection index at an intensity of 10 percent. The solid black line is the regression line. The dashed black lines indicate the trait means for GY and DTHD. The dashed red lines indicate the response to selection in both traits.

GY predictions in the TST set for HTP selection, univariate GS, and multi-trait GS were used with observed iid BLUPs for DTHD in the SP and index weights to develop the index predictions. The predictive abilities for HTP selection and univariate GS were similar in 2016-17, while in 2017-18 the predictive ability of univariate GS exceeded that of HTP selection (Fig. 6). As with predictions of yield directly, multi-trait GS showed the highest predictive abilities for both breeding cycles.

**Figure 6.**
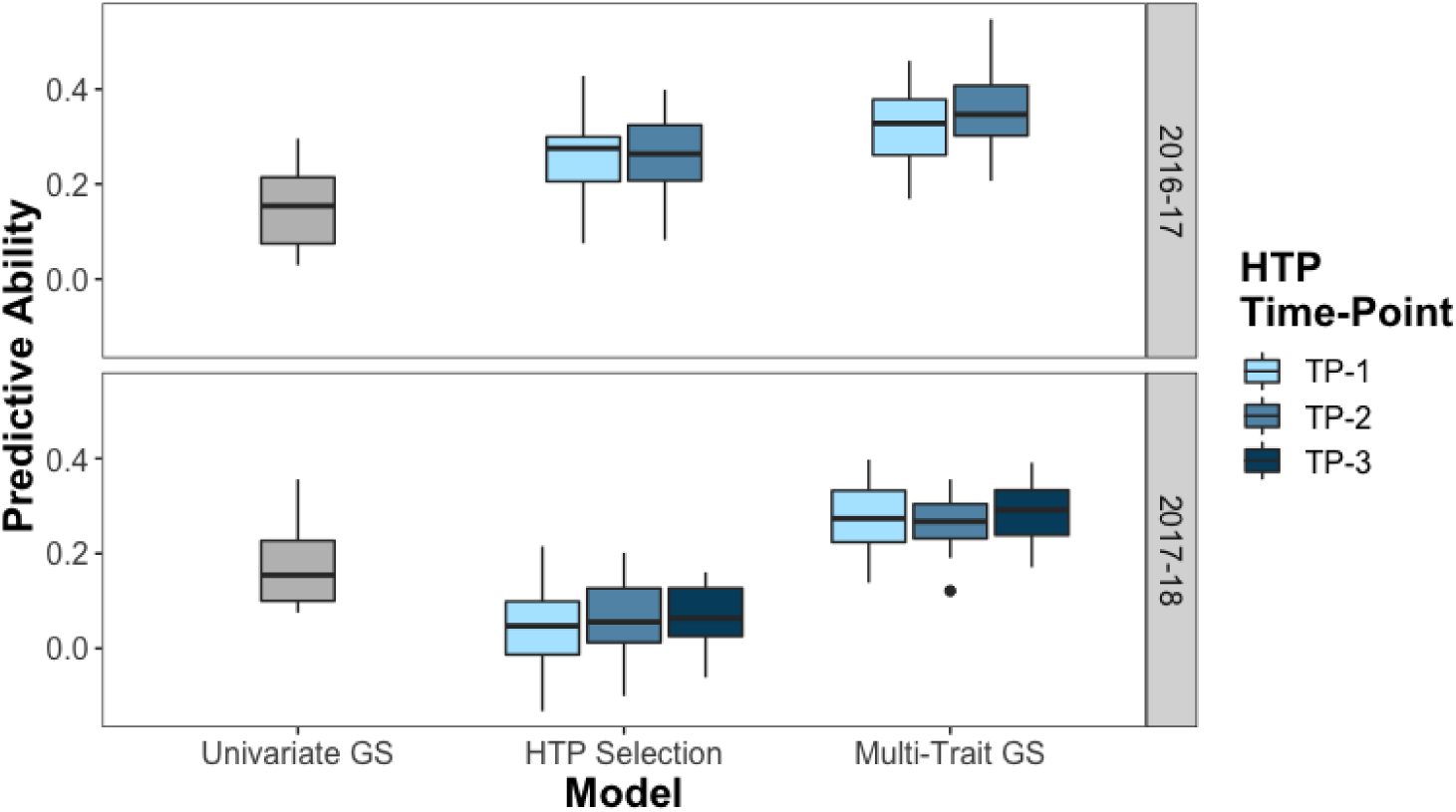
Predictive abilities of univariate GS, HTP selection, and multi-trait GS for a restricted selection index to increase GY while holding DTHD constant Predicted index values for the TST set were calculated using the predicted values for GY and observed iid BLUPs for DTHD in the SP. The predictive ability is expressed as the Pearson’s correlations between predicted values and observed index values calculated using the observed iid BLUPs for GY and DTHD in the YT across 20 random TRN-TST iterations. The univariate and multi-trait GS models were fit using the genomic relationship matrix (**G**).

## DISCUSSION

The aerial VI traits were found to be more heritable than GY in both the YT and SP, and minimal differences were observed with respect to heritability among the multiple time-points on which the HTP traits were recorded. Several previous studies have found aerial VI traits to be heritable and repeatable for wheat (Rutkoski et al., 2014; Crain et al., 2017; Sun et al., 2017), though the traits measured in these works were evaluated on replicated trials with plot sizes sufficient for reliably assessing yield. To our knowledge, this is the first report aiming to assess the heritability of aerial VI traits measured on small, unreplicated plots and estimate the extent to which they are predictive of GY in replicated yield trials.

It is possible that the estimates of narrow-sense heritability for traits measured in the SP are partially inflated. At the early stages of small grains breeding programs, breeders often sow full-sibs derived from the same set of parents adjacent to one another in the field for logistical ease and to enable visual comparisons among lines within families. The disadvantage of this approach, however, is that the genetic and spatial components of the phenotypic variance are confounded. When planting siblings together in families, Magnussen (1993) showed that the estimates of additive genetic variance and narrow-sense heritability were biased upward with positive spatial autocorrelation among neighboring plots. In this study, despite the possible confounding of genetic and spatial variance, the phenotypic and genetic correlations between the VI traits measured in the SP and GY in the YT were moderate. The second replicate of the YT was randomized, and the YT and SP also represented separate field experiments, each with independent environmental effects. In addition, the HTP traits in the SP_BP_ were also correlated with GY in the YT, despite having been measured during the prior breeding cycle and on smaller plot sizes than the SP. Together, these results suggest that the VI traits measured in at the early-generation seed-limited stages may still be effective for improving GY at the YT stage even if the genetic and spatial variance components are confounded as a result of planting full-sibs adjacent to one another.

Harvesting 300 plots of the SP_BP_ for GY enabled a direct comparison between HTP and CIMMYT’s visual selection scheme because all breeding lines, both selected and unselected by visual selection, were assessed for both GY and HTP. Although correlations between NDVI and GY in the SP_BP_ were weak, the NDVI traits were successful in identifying breeding lines with the lowest GY values for culling. The response to selection for GY was substantially greater using aerial NDVI as the selection criterion when compared to the current visual selection method. While these results are encouraging, the sample size of this experiment was relatively small. In addition, the GY values were based on the harvest of unreplicated 0.8m × 0.8m plots, and results from the YT and SP experiments showed that GY recorded in the SP is only moderately predictive of GY at the YT stage. Empirical validation is needed to confirm these results when GY is measured in replicated yield trials.

HTP traits have been shown to improve the ability to predict GY in wheat through multi-trait GS (Rutkoski et al., 2016; Sun et al., 2017; Crain et al., 2018); however, the ability to apply this approach to the seed-limited stage of breeding programs has not been previously evaluated. In this study, GY and HTP records assessed in the YT were integrated with genomic information to train multi-trait GS models. These were then applied to predict GY for the SP, where collecting HTP traits is possible but GY is not typically recorded. Overall, the multi-trait models showed the highest predictive abilities as compared to univariate GS and prediction with the HTP traits alone, though the increase over the latter was marginal. Therefore, it may be more cost-effective for breeding programs to invest in HTP alone as opposed to both HTP and genotyping for GS when selecting at the early-generation seed-limited stages. While the cost of genotyping has reduced greatly over the past decade, the large number of lines in breeding programs at the early stages can make applying GS expensive. However, if both genomic markers and HTP traits are available, using both sources of information to build prediction models would be optimal.

Multiple findings from this study suggested that using NDVI as a selection criterion at early breeding stages would have driven marked changes in DTHD within the breeding population. Mason and Singh (2014) observed similar directional changes in phenology when selecting on canopy temperature for improving wheat GY under drought. The potential for undesired directional response to selection in confounded traits is not unique to HTP but is rather relevant to any selection scheme in which other traits are confounded with the trait of interest. To account for the confounding effects of phenology, GY and NDVI traits were corrected for phenology by including DTHD as a fixed effect covariate during prediction. Given that strong correlations were observed between GY, DTHD, and NDVI, the predictive abilities of HTP selection and the multi-trait GS models were greatly reduced after accounting for DTHD to the extent that the addition of NDVI traits in multi-trait GS provided no additional predictive power compared to the univariate GS. Estimates of response to selection for GY in the YT were low, less than 1 percent for most NDVI time-points, when using DTHD-corrected NDVI traits evaluated in the SP as the selection criteria.

A limitation of this approach of accounting for trait confounding is that the genetic correlations of DTHD with GY and NDVI are ignored. Any of the genetic variance that may have been contributing to both traits is partitioned into the fixed effect. Selection indices, which take into account genetic correlations between traits, may therefore be a more suitable approach when traits are highly related. In this study, visual selections made in the SP_BP_ showed that the breeders are simultaneously accounting for relative maturity and do not select directionally on DTHD. A restricted selection index can mimic this by enabling selection for increased GY while holding DTHD fixed. This approach was utilized to compare the predictive abilities of HTP selection, univariate GS, and multi-trait GS to identify breeding lines with high GY potential across the distribution of DTHD. The results from 2016-17 showed that HTP selection gave similar levels of predictive ability as univariate GS when predicting the index, and that the multi-trait GS models that combined genomic marker information with HTP traits further increased predictive abilities. However, in 2017-18, HTP selection performed more poorly than univariate GS, and multi-trait models were only marginal better than the univariate case. These differences in predictive abilities were likely due to the index showing stronger correlations with VI traits in the SP in 2016-17 than in 2017-18. Rutkoski et al. (2016) showed that the level of gain in GS predictive ability from the addition of secondary traits is significantly associated with the genetic correlation between the secondary and target traits.

While VIs alone may not show a sufficiently strong and consistent genetic correlation with a restricted selection index for GY and DTHD to provide significant increases in GS predictive ability, it is likely that breeders will have access to more extensive suites of phenotypic traits in addition to VIs as HTP systems become increasingly automated, customizable, and scalable. For example, hyperspectral reflectance phenotypes, which record spectral reflectance at a large range of wavelengths, were shown to increase prediction accuracy for GY when compared to VIs (Montesinos-López et al., 2017). Beyond spectral reflectance-related traits, deep learning algorithms have been developed to determine DTHD from proximal imagery of wheat canopies (Wang et al., 2019), UAV imagery has been used to estimate lodging in wheat as a function of changes in plant height throughout the growing season (Singh et al., 2019), and convolutional neural networks have been trained to identify foliar diseases in maize from aerial imagery (Wu et al, 2019). Looking ahead, these comprehensive suites of traits may allow confounding effects such as those observed in this study to be adequately addressed through integrative selection approaches that account for genetic correlations between traits.

Until that time, the most practical approach may be to leverage useful information from HTP traits without relying on them fully. For example, van Ginkel et al. (2008) compared elite wheat breeding lines for GY under drought and heat stress developed through visual selection and through an integrated approach combining visual selection with measurements of canopy temperature depressions. Although the lines with the highest GY for both methods were statistically, similar, the integrated approach was more effective in identifying and eliminating the lower yielding lines. In this study, we showed that the VI traits could be used in conjunction with qualitative DTHD scores to identify breeding lines with higher GY potential for the “early”, “mid”, and “late” heading subgroups.

Although recent advancements in remote sensing and robotic technologies have produced inexpensive cameras, sensors, UAV, smartphone applications, etc., the overall cost of deploying HTP platforms can be considerable (reviewed by Reynolds et al., 2019). Additional expenses may include the labor required to utilize HTP platforms in the field, training of personnel and dedicated bioinformaticians, the development and implementation of data extraction pipelines, propriety software, and the establishment and maintenance of systems for storing and organizing large volumes of phenotypic data. Future improvements in the automation of HTP platforms and data analysis workflows as well as the development of open source software may help to alleviate some of the costs of implementing HTP, though they are not likely to decrease below the costs associated with visual selection. Therefore, the breeder must determine if the added investment in HTP can be offset by increases in the rate of genetic gain and/or the costs saved from sowing fewer replicated yield trials. Further studies are needed to compare empirical gains from selection per unit of cost using HTP versus current visual selection methods.

## ACKNOWLEDGEMENTS

This work was supported through the US Agency for International Development (USAID) Feed the Future Innovation Lab for Applied Wheat Genomics (Cooperative Agreement No. AID-OAA-A-13-00051). It was also supported in part by the Delivering Genetic Gain in Wheat project, supported by aid from the U.K. Government’s Department of International Development (DFID) and the Bill & Melinda Gates Foundation (Grant No.: OPP1133199). We are thankful to the National Science Foundation Graduate Research Fellowship (Grant No. DGE-1650441) and the U.S. Borlaug Fellows in Global Food Security program for supporting the graduate studies of Margaret R. Krause.

